# ca-circSCN8A promotes PASMCs ferroptosis via LLPS initiated R-Loop

**DOI:** 10.1101/2025.01.21.634205

**Authors:** Mengnan Li, Yingying Hao, Xinyue Song, Huiyu Liu, Chi Zhang, Jiaqi Zhang, Hanliang Sun, Xiaodong Zheng, Lixin Zhang, Hang Yu, Cui Ma, Xijuan Zhao, Daling Zhu

## Abstract

**BACKGROUND:** Ferroptosis has been implicated in pulmonary hypertension. Chromatin-associated RNAs are linked to ferroptosis. However, their role in pulmonary vascular ferroptosis in PH is unexplored.

**METHODS:** Bioinformatics, Sanger sequencing, RNase R and others were employed to identify differentially higher expression of ca-circSCN8A. Functional gain and loss assays were employed to unveil the role of ca-circSCN8A in hypoxia-induced redox-dependent ferroptosis in HPASMCs and PH mice model. Interaction between ca-circSCN8A and FUS was detected via RNA immunoprecipitation and pull-down assays. FRAP, CHIRP-qPCR, MDA, GSH and GSSG were conducted to explore the potential molecular mechanism.

**RESULTS:** ca-circSCN8A was firstly identified and confirmed to be upregulated in PH. Upregulation of ca-circSCN8A can promote hypoxia induced ferroptosis in HPASMCs. Under hypoxic conditions, ca-circSCN8A promoted the lactylation of FUS by recruiting EP300, leading to formation of LLPS of ca-circSCN8A/FUS/EP300. LLPS maintained the stability of the R-Loop formed by ca-circSCN8A and ferroptosis related-gene SLC7A11 promoter that inhibit the transcription of the SLC7A11 gene, further result in the disruption of the redox balance and causes ferroptosis in HPASMCs.

**CONCLUSIONS:** ca-circSCN8A recruits EP300 to promote the lactylation of FUS, thereby promoting LLPS formation of ca-circSCN8A/EP300/FUS. Further, the LLPS motivates ca-circSCN8A to form R-loop with its non-host SLC7A11 and participating in the regulation of hypoxic induced HPASMCs redox-dependent ferroptosis. This is the first confirmation that circRNAs forming R-loops with non-host genes regulated by LLPS. Our findings indicated that ca-circSCN8A plays an important role in ferroptosis of hypoxic induced HPASMCs and may be a potential target for the treatment of PH.

## INTRODUCTION

Pulmonary hypertension (PH) is a life-threatening cardiovascular disease.^1^ Pulmonary vascular remodeling (PVR) is the key pathological change in PH,^2,3^ involving endothelial cells (PAECs), smooth muscle cells (PASMCs), fibroblasts, and inflammatory cells.^4–6^ Various cellular pathophysiological changes contribute to PVR, including cell proliferation, programmed cell death, and autophagy. Ferroptosis, a form of programmed cell death distinct from apoptosis, necrosis, and autophagy, is characterized by iron-dependent lipid peroxidation.^7^ It is regulated by oxidative– reductive homeostasis, iron metabolism, mitochondrial activity, amino acid/lipid/glucose metabolism, and related signaling pathways. Although previous studies have described ferroptosis in PH-related cells, the mechanisms involved remain unexplored.

Transcriptional abnormalities in ferroptosis-related genes represent initial and pivotal factors.^8^ The transcription of several genes is altered in ferroptosis-prone cells. For example, the expression levels of COX2, ACSL4, and NOX1 are increased, whereas those of GPX4, SLC7A11, and ferritin are decreased.^9^ The transporter SLC7A11, which facilitates the exchange of cysteine and glutamate, plays a crucial role in preserving cellular redox homeostasis and preventing ferroptosis.^10, 11^ Multiple mechanisms, including transcriptional regulation, epigenetic control, and posttranscriptional regulation, govern SLC7A11 mRNA levels, protein stability, subcellular localization, and transporter activity and may be involved in mediating ferroptosis.^12^ Given that transcription serves as the initial step of gene function, we focused on elucidating the specific regulatory factors and pathways that control SLC7A11 transcription.

During gene transcription, newly synthesized RNA can base pair with open double-stranded DNA, forming stable DNA‒RNA–DNA triplex structures (known as R-loops).^14^ Abnormal R-loop accumulation can cause conflicts between transcription and replication machines, leading to genomic instability. Thus, R-loops play a crucial role in gene transcription.^15^ Chromatin-associated circular RNAs (ca-circRNAs) are a category of circular RNA molecules that interact with chromatin, influencing gene expression, chromatin structure, and cellular function.^16–18^ CircRNAs have been reported to positively or negatively regulate the transcription of their host genes by binding to RNA polymerase II (Pol II), recruiting proteins or forming an R-loop to target transcriptional regulatory regions.^19^ While current research has focused mainly on circRNA-mediated R-loop formation in host gene transcription, whether ca-circRNA can regulate transcription by forming R-loops with nonhost genes remains unknown.^20^ In addition, it remains unclear whether ca-circRNA maintains the stability of promoter-associated R-loops through protein complexes. Therefore, elucidating the protein composition involved in the R-loop formation of ca-circRNA and nonhost gene promoters, particularly the stability of the complex maintaining promoter-associated R-loops, will provide critical insight into gene transcription regulation.

Liquid‒liquid phase separation (LLPS) drives the formation of membraneless cellular organelles by large biomolecules, such as proteins and RNA. LLPS plays a crucial role in physiological and pathological processes^21,22^ and is essential for transcriptional regulation.^23^ LLPS selectively enriches or excludes relevant molecules, increasing the probability of molecular interactions and altering molecular conformations, facilitating or inhibiting biochemical reactions.^24^ As promoter-related R-loops occur at the proximal pause site, which is always less than 60 bp, their stability should be linked to protein complexes. Hence, LLPS may serve as a mechanism that facilitates the formation of R-loops between ca-circRNAs and nonhost genes, further regulating nonhost gene transcription.

This study aimed to investigate whether hypoxia induces the formation of an R-loop in HPASMCs, specifically involving the ca-circRNA and SLC7A11 gene promoter regions. Moreover, we explored whether the downregulation of SLC7A11 can lead to an imbalance in the oxidative-reductive state of HPASMCs, ultimately mediating ferroptosis.

## MATERIAL AND METHODS

The details of the methods and statistical analyses are provided in the Supporting Material.

## RESULTS

### The expression and subcellular distribution of ca-circSCN8A in hypoxic HPASMCs

A PH-related dataset (GSM5235379-GSM5235391) was selected from the GEO database. The five most significantly upregulated circRNAs were screened according to the following criteria: log2 (fold change>| 2 |) and P<0.01. The qRT‒PCR results indicated that hypoxia significantly increased the expression of ca-circSCN8A and circWNT5B in HPASMCs (Figure 1A and B). Sanger sequencing was performed on the PCR products, and the results revealed that only ca-cicSCN8A was connected at the head and tail, confirming its circular RNA nature. Ca-circSCN8A is 853 nucleotides long and located at chr12:52180325–52188425. The cyclization site is formed by exons 21–25 (Figure 1C). Ca-cicSCN8A demonstrated stronger resistance to RNase R than did linear SCN8A (Figure 1D). Moreover, significant ca-circSCN8A expression upregulation was observed in HPASMCs compared with that in HPAECs (Figure 1E). A time point validation revealed that the most significant ca-circSCN8A expression upregulation in HPASMCs occurred after 24 h of hypoxia (Figure 1F). The results of the RNA FISH assay revealed that ca-circSCN8A expression was upregulated under hypoxic conditions compared with that in normoxia, and it was distributed in both the cytoplasm and nucleus (Figure 1G). The tissue-specific expression of ca-circSCN8A in hypoxic and normoxic lung tissues was analyzed. The expression of ca-circSCN8A in hypoxic lung tissue was significantly greater than that in normoxic lung tissue (Figure 1H). The distribution of ca-circSCN8A in lung tissue was determined by FISH, and it was found to be expressed mainly in pulmonary arteries (Figure 1I).

**Figure 1:**
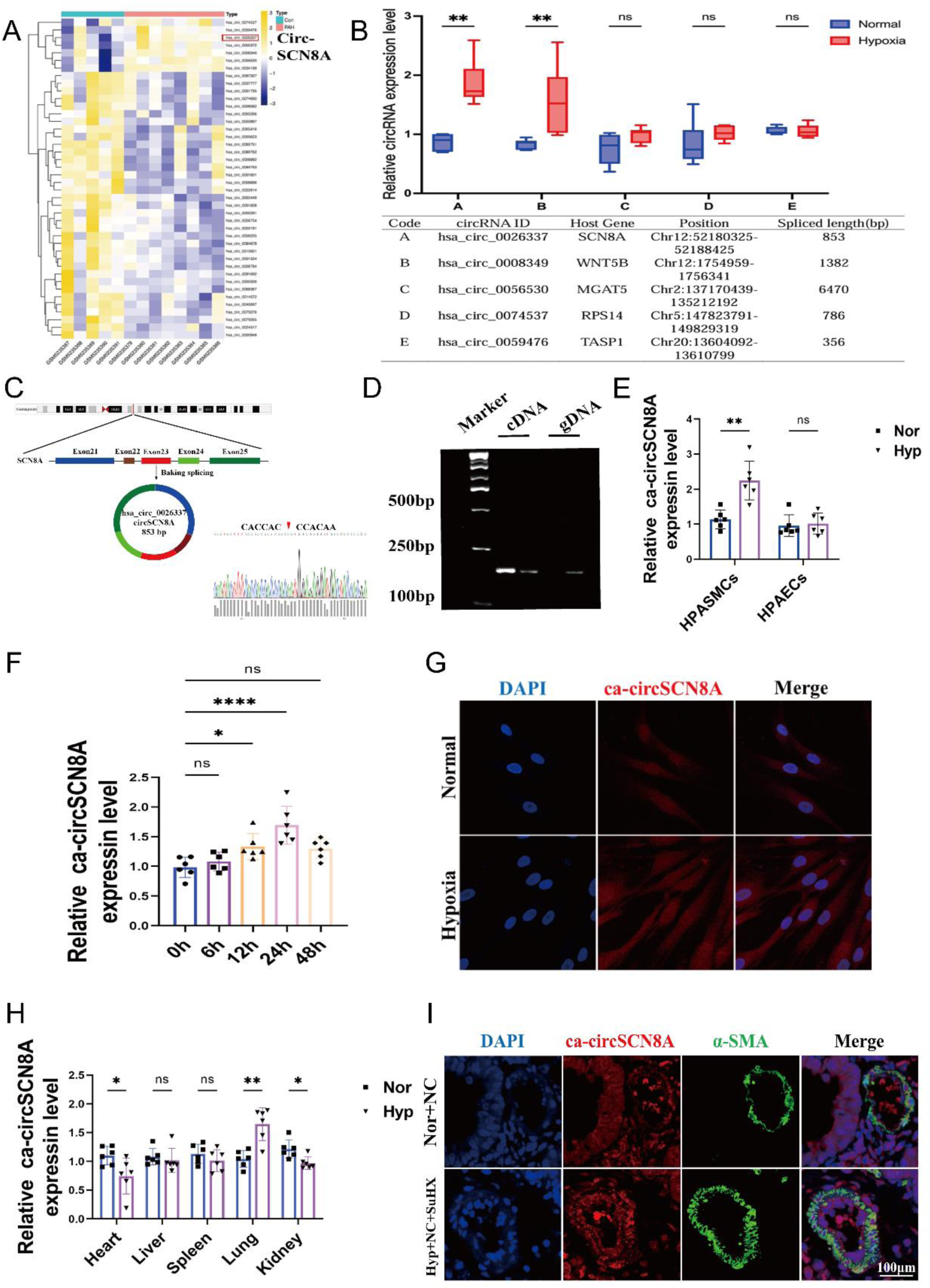
The effects of hypoxia on the expression and subcellular localization of ca-circSCN8A. (A) Expression profile of circRNAs in patients with PH in the GEO database. (B) Expression of five circRNA types under hypoxic conditions (n = 6). *P<0.05, **P<0.01 vs the normal group. (C) Genomic schematic of ca-circSCN8A and Sanger sequencing results. (D) qRT‒PCR products with divergent and convergent primers showing circularization of ca-circSCN8A (cDNA [complementary DNA], gDNA [genomic DNA]; n = 6). (E) ca-circSCN8A expression in HPAECs and HPASMCs (n = 6). (F) Ca-circSCN8A expression at different time points in hypoxic HPASMCs (n = 6). (G) RNA FISH results of ca-circSCN8A expression and subcellular localization in HPASMCs exposed to normoxia or hypoxia for 24 h. ca-circSCN8A (red) and DAPI (blue nuclear) were used. Scale bar = 200 µm. (H) Ca-circSCN8A expression in the heart, liver, spleen, lung, and kidney (n = 6). (I) FISH results of ca-circSCN8A distribution in lung tissue. The data are presented as the means ± SDs. Statistical analyses were performed by 1-way ANOVA followed by Bonferroni correction and Student’s t test for 2 means. *P<0.05, **P<0.01, ***P<0.001.

### Upregulated ca-circSCN8A promotes PVR and increases ferroptosis

The role of ca-circSCN8A in hypoxia-induced PH was investigated in vivo. A smooth muscle-specific ca-circSCN8A knockout mouse model with hypoxic PH was constructed via AAV9 ca-circSCN8A, and the efficiency of the knockdown was validated (Figure 2A). Right ventricular systolic pressure (Figure 2B) and the right ventricular hypertrophy index (Figure 2C) were measured, revealing a significant reversal of these pH indicators when ca-circSCN8A was inhibited. Echocardiography measurements suggested that the decrease in the pulmonary artery velocity time integral and pulmonary artery acceleration time caused by hypoxia could be attenuated by silencing ca-circSCN8A (Figure 2D). Analyses of pulmonary artery morphology and pulmonary vascular thickness through HE (Figure 2E), α-SMA (Figure 2F), and Masson (Figure S2A) staining indicated that distal PVR induced by hypoxia was reversed by knockdown of ca-circSCN8A. The expression levels of the ferroptosis marker ACSL4 were increased by hypoxia, whereas those of SLC7A11 and GPX4 were decreased. These expression changes could be reversed by knocking down ca-circSCN8A under the same conditions (Figure S2B).

A ca-circSCN8A overexpression plasmid was constructed and transfected into HPASMCs, and the overexpression efficiency was verified (Figure S2C). The cells were cocultured with an apoptosis inhibitor (ZVAD-FMK), necrosis inhibitor (Nec-1), or ferroptosis inhibitor (Fer-1) under normoxic conditions. The cell death rate caused by the overexpression of ca-circSCN8A was significantly reversed only by Fer-1 (Figure S2D). To further explore whether hypoxia regulates HPASMC ferroptosis through ca-circSCN8A, ca-circSCN8A antisense oligonucleotides (ASO-ca-circSCN8A) were used to knockdown ca-circSCN8A, the knockdown efficiency was verified, and ASO-001 was selected for subsequent experiments (Figure S2E). The increase in the ferroptosis-related redox homeostasis indicators MDA, Fe^2+^, and GSH and the decrease in GSSG caused by hypoxia were significantly reversed by the knockdown of ca-circSCN8A (Figure 2G). Similarly, electron microscopy revealed that mitochondrial morphology significantly recovered after knocking down ca-circSCN8A (Figure S2F). WB results revealed that the expression of the ferroptosis marker ACSL4 in HAPSMCs was increased by hypoxia, whereas the expression of SLC7A11 and GPX4 was decreased. These expression changes could be reversed by knocking down ca-circSCN8A under the same conditions (Figure 2H). A single-factor functional recovery experiment revealed that the overexpression of ca-circSCN8A promoted HPASMC ferroptosis under normoxic conditions (Figure S2G–I). These results indicate that ca-circSCN8A plays a key role in promoting hypoxia-related HPASMC ferroptosis and PVR.

**Figure 2:**
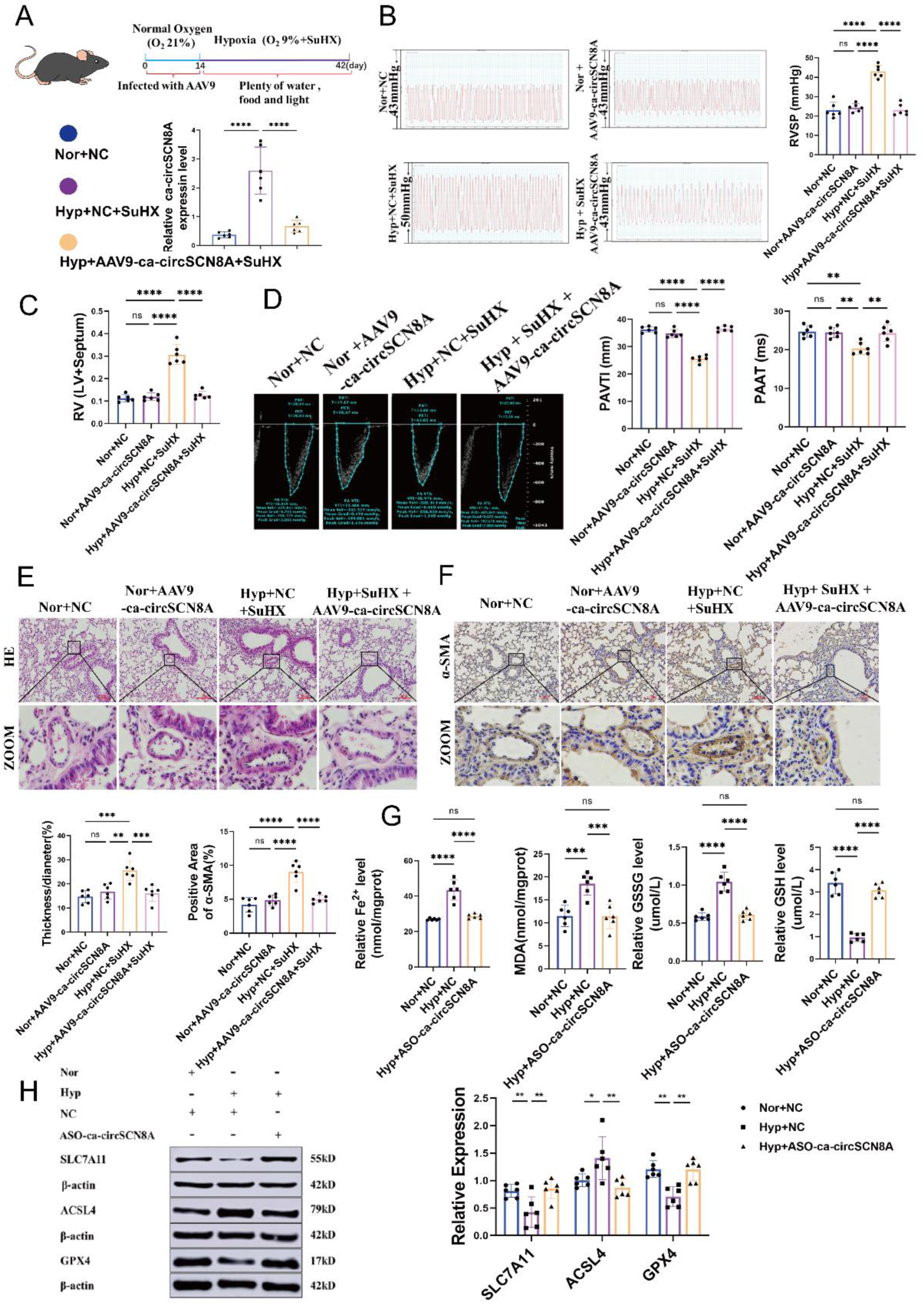
Hypoxia promotes PVR and ferroptosis via ca-circSCN8A. (A) Construction of a mouse AAV9 plasmid on the basis of the target sequence. The mice were infected with AAV9 once under normal hypoxic conditions for 14 days, with HYP lasting for 28 days. (B) Right ventricular systolic pressure (RVSP) in the hypoxia-induced PH mouse model and (C) the weight ratio of the right ventricle (RV) to the left ventricle (LV) + septum (S) (n = 6). (D) Pulmonary artery velocity time integral (PAVTI) and pulmonary artery acceleration time (PAAT) in serum 9-type adenovirus-related virus (AAV9)-ca-circSCN8A-treated mice (n = 6). (E) Pulmonary artery morphology analysis was performed via hematoxylin and eosin (HE) staining, and (F) α-smooth muscle actin (α-SMA) staining was used to measure pulmonary vascular thickness (n = 6). Scale bar = 100 µm. All values are expressed as the means ± SEMs. (G) Effects of ca-circSCN8A on MDA, Fe^2+^, GSSG, and GSH expression (n = 6). (H) WB results showing the effects of ca-circSCN8A on the expression of the ferroptosis-related proteins ACSL4, SCL7A11, and GPX4 in HPASMCs (n = 6).

### ca-circSCN8A mediates ferroptosis in HPASMCs via SLC7A11

A ceRNA network functional enrichment analysis was conducted to elucidate how hypoxia-induced ferroptosis in HPASMCs is mediated by ca-circSCN8A. The results revealed a significant association with the regulation of gene expression (Figure 3A). qRT‒PCR demonstrated that hypoxia modulates the expression of SLC7A11 via ca-circSCN8A (Figure 3B). ASO-ca-circSCN8A and si-SLC7A11 were cotransfected to ascertain whether ca-circSCN8A mediates ferroptosis in HPASMCs through SLC7A11 by measuring ferroptosis-related indicators. si-SLC7A11 reversed the alterations in Fe^2+^, MDA, GSSG, and GSH levels (Figure 3C and D). WB revealed that si-SLC7A11 could reverse the changes in ACSL4 and GPX4 expression (Figure 3E). Collectively, these findings suggest that ca-circSCN8A mediates ferroptosis in HPASMCs by modulating SLC7A11 expression.

### SLC7A11 expression is regulated by the formation of an R-loop between ca-circSCN8A and the SLC7A11 promoter

Sequence alignment was performed to further explore how ca-circSCN8A regulates the mRNA level of SLC7A11, revealing that two binding motifs are shared between ca-circSCN8A and the SLC7A11 promotor (Figure 3F). In addition, the R-Loop Base program indicated that these motifs are part of the R-loop-forming sequence of SLC7A11 and are located at chromatin chr4:138209522–138211044 (ca-circSCN8A127–136, SLC7A11 274–283) and chr4:138153055–138242533 (ca-circSCN8A 95–105, SLC7A11 1460–1470) (Figure 3G). Therefore, we hypothesized that ca-circSCN8A may regulate SLC7A11 expression by forming an R-loop with the SLC7A11 gene. To confirm this hypothesis, we first examined the expression levels of R-loops in HPASMCs under normoxic and hypoxic conditions. Our results revealed increased expression of R-Loop in HPASMCs under hypoxic conditions (Figure 3H). Moreover, the overexpression of RNase H1 in hypoxic HPASMCs, which was intended to reduce R-loop levels, significantly reversed the hypoxia-induced decrease in SLC7A11 mRNA expression (Figure 3I). A biotin probe that complements the reverse sequence of ca-circSCN8A was designed to determine whether the elevated R-loop is formed by the interaction between ca-circSCN8A and the SLC7A11 promoter. ChIRP-qPCR experiments further confirmed that ca-circSCN8A forms an R-loop with the SLC7A11 promoter of the ferroptosis gene via site 1 and that the R-loop increases significantly under hypoxic conditions (Figure 3J). Upon mutation of site 1, the hypoxia-induced alterations in Fe^2+^, MDA, GSSG, and GSH (Figure S3A–D) and the changes in SLC7A11 (Figure 3K), ACSL4, and GPX4 (Figure 3L) protein expression were reversed. These findings underscore the role of R-loop formation between ca-circSCN8A and the SLC7A11 gene promoter in mediating the ferroptosis of HPASMCs.

**Figure 3:**
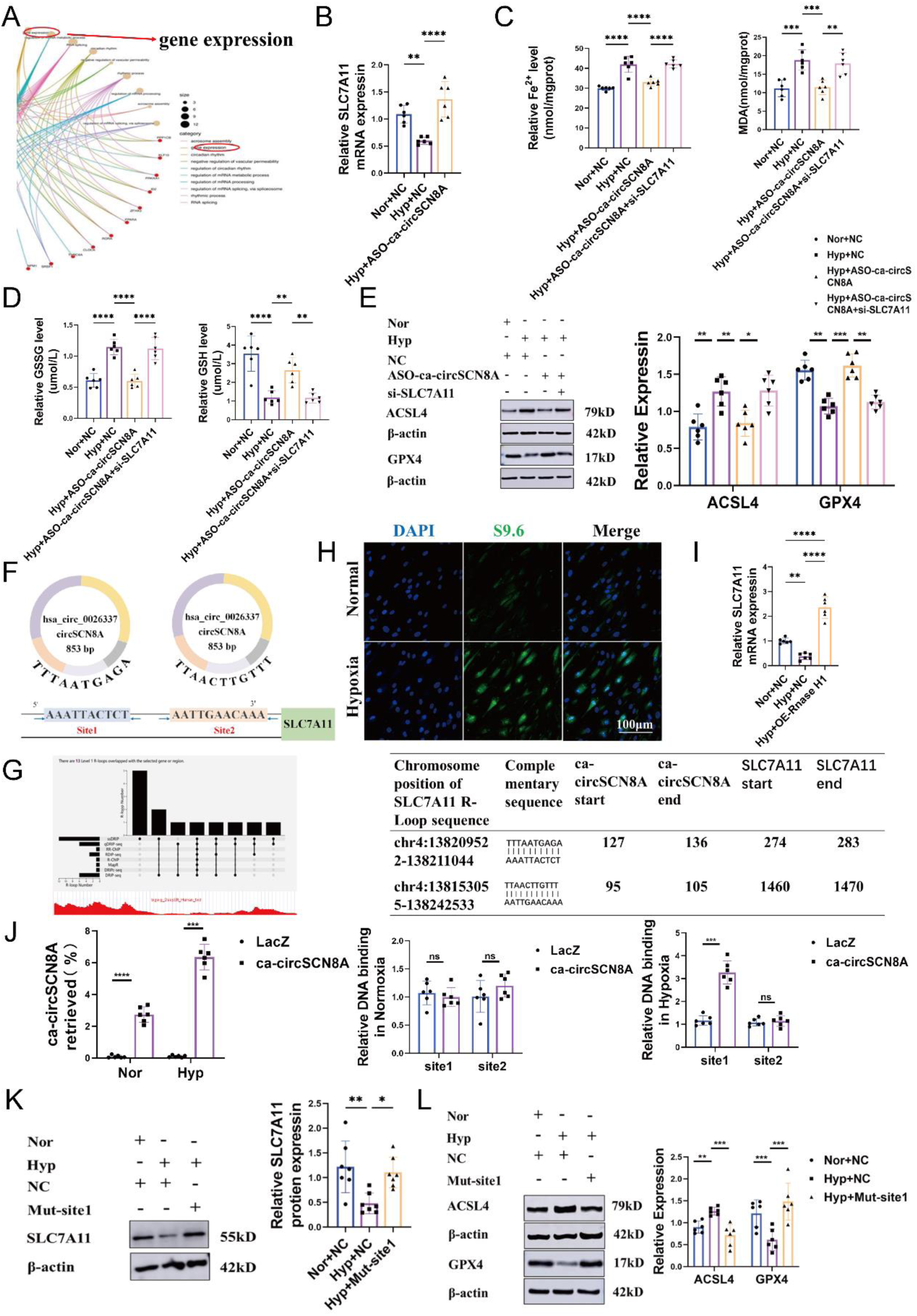
ca-circSCN8A mediates HPASMC ferroptosis by forming an R-loop with the SLC7A11 gene promoter. (A) ceRNA network enrichment analysis. (B) SLC7A11 mRNA expression. (C/D) MDA, Fe^2+^, GSH, and GSSG expression levels (n = 6). (E) WB results showing the expression of the ferroptosis markers ACSL4 and GPX4 (n = 6). (F) Immunofluorescence images of R-Loop in HPAMSCs under normoxic and hypoxic conditions. (G) SLC7A11 mRNA expression after transfection with RNase H1 (n = 6). (H) Prediction of the formation site of the SLC7A11 R-loop and SSDRIP-seq. (I) Sequence alignment analysis and related information for ca-circSCN8A and SLC7A11. (J) ChIRP-qPCR results verifying the formation of an R-loop between the ca-circSCN8A promoter and the SLC7A11 promoter (n = 6). (K/L) SLC7A11, ACSL4, and GPX4 expression in normoxic, hypoxic, and site 1 mutant plasmid-treated HPAMSCs (n = 6).

### R-Loop formation is regulated by the ca-circSCN8A downstream protein FUS

The RBP suit and RBPDB programs were employed to predict downstream binding proteins of ca-circSCN8A (Figure 4A). Proteins with a perfect score of 100% were selected for molecular docking. FUS demonstrated the highest binding affinity with ca-circSCN8A, with a binding energy of –8.4 kcal/mol (Figure 4B). Consequently, we targeted the DNA/RNA-binding protein FUS for subsequent exploration. RNA pull-down (Figure 4C) and RIP (Figure 4D) assays further confirmed the interaction between FUS and ca-circSCN8A in HPASMCs. The linear SCN8A probe was confirmed not to interact with FUS (Figure S4A and B). The catRAPID program indicated that their binding region was 226–277 nt (Figure 4E). After segmenting ca-circSCN8A and reperforming the RNA pull-down assay, FUS was detected only in the complex pulled down with the 226–277 nt sequence. These findings further suggest that ca-circSCN8A interacts with FUS via the 226–277 region (Figure 4F). The FISH-IF results revealed colocalization of ca-circSCN8A and FUS within the cell nucleus (Figure 4G).

We subsequently validated the function of FUS in HPASMCs. WB revealed hypoxia-induced upregulation of FUS expression (Figure 4H). After constructing interference fragments of FUS and confirming their interference efficiency (Figure S4C), we selected si-FUS-181 for further investigation. Immunofluorescence results revealed that si-FUS-181 significantly reversed the hypoxia-induced increase in R-Loop formation (Figure S4D). Moreover, when FUS was knocked down, R-loop formation was significantly reduced (Figure S4E). The qRT‒PCR and WB results from the same batch of cell samples indicated that FUS significantly reversed hypoxia-induced changes in SLC7A11 protein and mRNA levels (Figure 4I), suggesting that hypoxia regulates SLC7A11 protein and mRNA expression via FUS. Concurrently, FUS knockdown significantly reversed the changes in the expression of ferroptosis markers (Figure S4F–H). These findings indicate that hypoxia regulates SLC7A11 expression and HPASMC ferroptosis via FUS. A FUS overexpression plasmid was constructed to explore whether FUS is required for R-loop formation between ca-circSCN8A and the SLC7A11 promoter. After verifying the overexpression efficiency (Figure S4I), we cotransfected ASO-ca-circSCN8A and OE-FUS into HPASMCs and assessed R-loop and SLC7A11 expression levels and ferroptosis-related indicators. Immunofluorescence analysis revealed that OE-FUS significantly reversed the R-loop expression caused by ca-circSCN8A (Figure 4J). The qRT‒PCR and WB results from the same batch of cell samples suggested that OE-FUS significantly reversed the hypoxia ASO-ca-circSCN8A-induced changes in SLC7A11 protein and mRNA expression (Figure S4J) and ferroptosis-related indicators (Figures 4K and S4K). It can be concluded that ca-circSCN8A regulates the expression of SLC7A11 and the ferroptosis of HPASMCs through its downstream DNA/RNA binding protein FUS, forming an R-loop with the SLC7A11 promoter.

**Figure 4:**
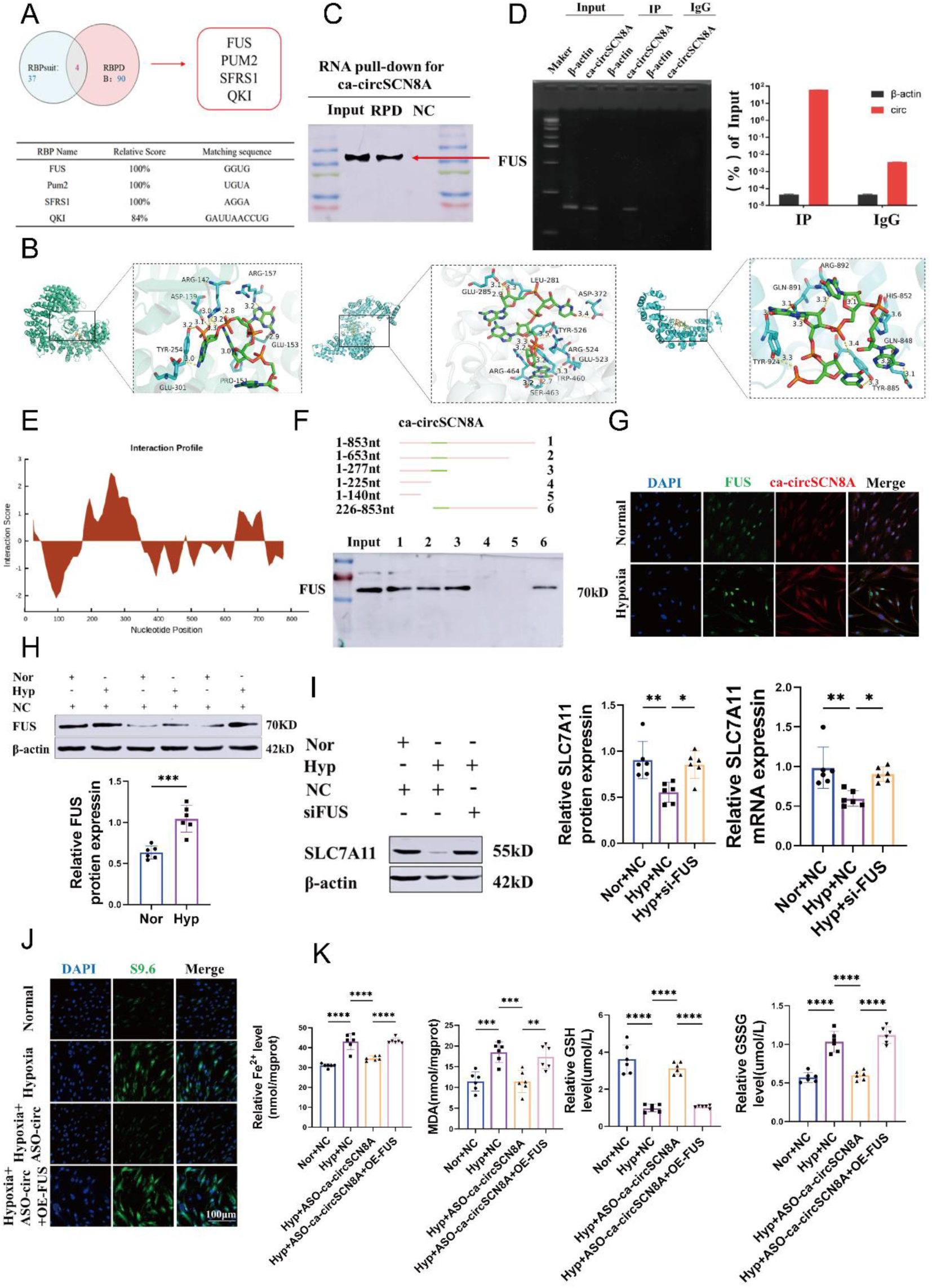
SLC7A11 expression is regulated by ca-circSCN8A via its downstream DNA/RNA binding protein FUS. (A) Downstream binding proteins of ca-circSCN8A were predicted by RBPsuit and RBPDB. (B) Three-dimensional molecular docking results of ca-circSCN8A with FUS/PUM2/SFRS1 were predicted and visualized via the HNADOCK server. (C) Protein complexes pulled down by ca-circSCN8A were identified, followed by immunoblot analysis of FUS after the pull-down experiment (n = 3). (D) The correlation between FUS and ca-circSCN8A expression was determined via RNA immunoprecipitation (RIP) (n = 3). (E) The binding region of ca-circSCN8A with FUS was predicted by catRAPID. (F) ca-circSCN8A is divided into six segments (1– 6), and its binding sequence with FUS is visualized. The interaction between ca-circSCN8A and FUS from 226–277 nt was further confirmed by RNA pull-down experiments. (G) Colocalization of ca-circSCN8A (red) and FUS (green) in the cell nucleus was verified by FISH-IF. Scale bar = 10 μm. (H) FUS expression in HPASMCs under hypoxic and normoxic conditions was measured via WB. (I) SLC7A11 protein and mRNA expression levels were measured by WB and qRT‒PCR after transfection with si-FUS-181 (n = 6). (J) Immunofluorescence images of R-Loop in HPASMCs transfected with ASO-ca-circSCN8A and si-FUS-181. (K) The expression levels of Fe^2+^, MDA, GSH, and GSSG in HPASMCs under hypoxic and normoxic conditions were measured with the relevant reagent kits (n = 6). All values are presented as the means ± SEMs. ASO indicates antisense oligonucleotide. Nor, normoxia; Hyp, hypoxia.

### The formation of LLPS with FUS induced the R-loop formation

To further explore how FUS induced the formation of R-loops, the expression of FUS was initially examined in normoxic and hypoxic HPASMCs following ca-circSCN8A knockdown. The results indicated that FUS expression was not modulated by ca-circSCN8A (Figure 5A). Subsequent bioinformatics analysis of the cellular components of ca-circSCN8A revealed a strong correlation with nuclear specks (Figure 5B). Previous studies have demonstrated that FUS can undergo LLPS.^25^ Given the absence of prior reports of circRNAs forming R-loops with nonhost genes, the propensity of LLPS to selectively enrich or exclude related molecules, increase the likelihood of molecular interactions, and even alter molecular conformation to facilitate or inhibit biochemical reactions provides a plausible mechanism for ca-circRNA and nonhost genes to form R-loops and regulate the transcription of nonhost genes.^26^ On the basis of these findings, we hypothesized that ca-circSCN8A might suppress its expression by performing LLPS with FUS and binding to the R-loop of the SLC7A11 promoter. To test this hypothesis, we first observed nuclear condensates formed by ca-circSCN8A and FUS in the cell nuclei of normoxic and hypoxic HPASMCs. These nuclear condensates were significantly prevalent under hypoxic conditions (Figure S5A). Subsequently, green fluorescent protein (GFP)-fused FUS and mCherry-fused ca-circSCN8A expression vectors were constructed and transfected into HPASMCs. The results revealed that these proteins form nuclear condensates in living cells (Figure 5C). To verify the liquid properties of this nuclear condensate, droplet fusion experiments and fluorescence recovery experiments were performed after photobleaching. The results showed that small particles in HPASMCs can fuse into larger particles over time (Figure 5D). Molecules in the condensate can diffuse and exchange with the surrounding solution (Figure 5E), further substantiating the liquid-like properties of the condensate. These findings indicate that ca-circSCN8A and FUS undergo LLPS in hypoxic HPASMCs.

We subsequently explored the relationships among LLPS, SLC7A11 expression, and ferroptosis. HPASMCs were treated with the LLPS inhibitor 1,6-HD, and the expression of SLC7A11 and ferroptosis-related indicators was measured. The results suggested that 1,6-HD can significantly reverse the expression of SLC7A11 caused by hypoxia (Figure 5F). Concurrently, inhibiting LLPS significantly reversed the changes in the expression of ferroptosis markers (Figure 5G and H). These findings indicate that hypoxia regulates SLC7A11 expression and HPASMC ferroptosis through LLPS.

Finally, we further explored the relationship between LLPS and R-loop formation. FUS ChIP-seq suggested that FUS was significantly enriched in the gene promoter region (Figure 5I). FUS has inherently disordered regions (IDRs). IDRs regulate the biological function of R-loops by directly identifying the R-loop structure and recruiting other R-loop processing factors and LLPS-mediated R-loop structural assembly mechanisms. IP-IB results revealed that FUS only bound to the R-loop in hypoxic HPASMCs (Figure 5J). To detect the change in R-loop levels in hypoxic HPASMCs after FUS deletion, we introduced the FKBPDF36V degradation tag and used d-TAG to induce rapid depletion of endogenous FUS. We found that in hypoxic FUS-dTAG HPASMCs, the R-loop level was significantly reduced (Figure 5K). Endogenous depletion of FUS reversed the expression of SLC7A11 mRNA induced by hypoxia (Figure 5L). In addition, a dot blot analysis revealed that hypoxia increased the number of R-loops, and the LLPS inhibitor 1,6-HD reversed this phenomenon (Figure 5M). Moreover, 1,6-HD treatment of cells reversed the expression of SLC7A11 mRNA induced by hypoxia in a time-dependent manner (Figure 5N). These findings indicate that ca-circSCN8A suppresses SLC7A11 expression and ferroptosis by forming LLPS with FUS and binding to the R-loop of the SLC7A11 promoter.

**Figure 5:**
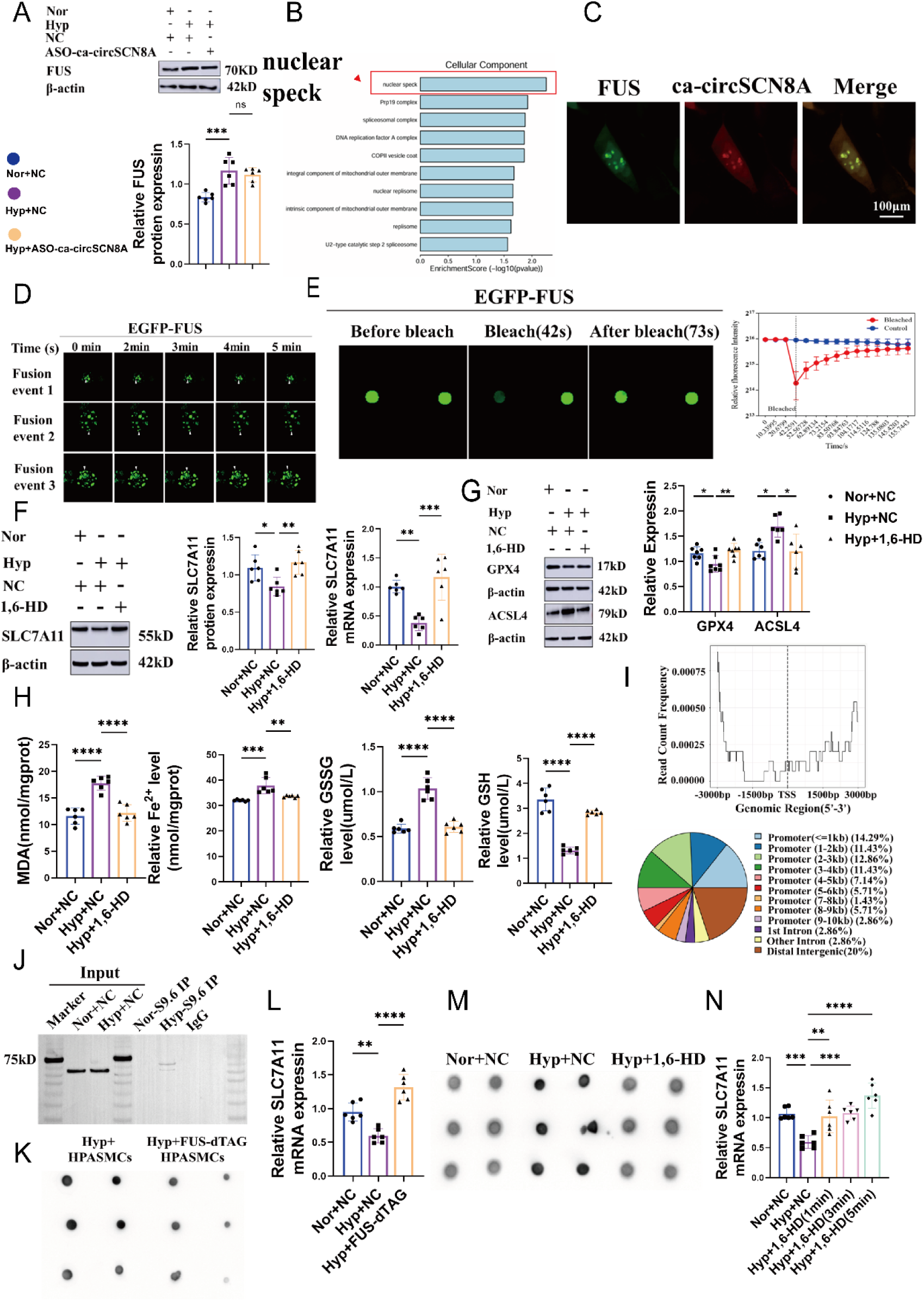
The expression and ferroptosis of SLC7A11 were inhibited by the LLPS structure formed by ca-circSCN8A binding with FUS and the SLC7A11 promoter R-loop. (A) FUS expression was detected via WB. (B) Bioinformatic analysis of the cellular components of ca-cicSCN8A. (C) Expression vectors of green fluorescent protein (GFP) fused with FUS and mCherry fused with ca-circSCN8A were transfected into living cells, and the formation and distribution of nuclear aggregates were observed via confocal microscopy. Scale bar = 100 μm. (D) A droplet fusion experiment was performed. (E) Fluorescence recovery after the photobleaching experiment was conducted. (F) The mRNA and protein levels of SLC7A11 were measured by qRT‒PCR and WB, respectively (n = 6). (G) The expression levels of the ferroptosis marker proteins ACSL4 and GPX4 were measured via WB. (H) The expression levels of Fe^2+^, MDA, GSH, and GSSG in hypoxic and normoxic HPASMCs were measured with related reagent kits (n = 6). (I) CHIP-seq data for FUS were analyzed. (J) The level of R-loop formation by FUS in normoxic and hypoxic HPASMCs was detected by IP–IB. (K) R-Loop formation in HPASMCs with endogenous depletion of FUS under hypoxic and hypoxic conditions was detected by dot blot. (L) Expression of SLC7A11 mRNA in HPASMCs with endogenous FUS deficiency under normoxic and hypoxic conditions. (M) R-loop expression in HPASMCs treated with 1,6-HD under normoxia, hypoxia, and hypoxia conditions was detected via dot blot. (N) SLC7A11 mRNA levels in HPASMCs with endogenous depletion of FUS under normoxic and hypoxic conditions and in HPASMCs treated with different concentrations of 1,6-HD under hypoxic conditions were detected via qRT‒PCR (n = 6).

### LLPS formation is regulated by the lactylation modification writer EP300

To reveal the mechanism behind the formation of LLPS, we employed the STRING database to predict proteins downstream of FUS. Our findings indicated tight binding between FUS and EP300 (Figure 6A). This interaction was further confirmed in HPASMCs through a co-IP experiment (Figure 6B). A subsequent protein‒protein interaction simulation suggested stable binding between FUS (gold) and EP300 (blue) via LYS510 (Figure 6C). DeepKla analysis revealed LYS510 as the lactylation site of FUS (Figure 6D). WB revealed the upregulation of EP300 expression under hypoxic conditions (Figure 6E). On the basis of these findings, we hypothesized that the LLPS formed by ca-circSCN8A and FUS could be regulated by the lactylation modification “writer” EP300. To verify this hypothesis, we first measured lactate levels in normoxic and hypoxic HPASMCs. The results revealed increased lactate levels in HPASMCs under hypoxia (Figure S6A). Coculturing hypoxic HPASMCs with 20 mmol of the lactate inhibitor DCA and 50 nmol of the lactate agonist ROT led to significant increases and decreases in the lactate and lactylation levels induced by hypoxia, respectively (Figure S6B). Concurrently, the number of nuclear aggregates formed by LLPS in hypoxic HPASMCs treated with DCA was significantly reduced (Figure S6C). DCA significantly reversed the expression of SLC7A11 induced by hypoxia (Figure S6D). Detection via related reagent kits and WB results revealed that DCA significantly reversed the changes in the expression of ferroptosis indicators (Figure S6E and S6F).

We subsequently constructed an EP300 interference fragment and verified its efficiency (Figure S6G). Upon transfection into hypoxic HPASMCs and detection of lactate expression levels, we found that knocking down EP300 could significantly reduce the increase in lactate expression caused by hypoxia (Figure 6F). Treating hypoxic HPASMCs with different concentrations of the lactate inhibitor C646 led to a concentration-dependent reduction in the increase in lactate expression caused by hypoxia (Figure S6H). Moreover, immunoprecipitation and immunoblotting experiments revealed a reduction in the lactylation level of FUS after EP300 was knocked down (Figure 6G). These results indicate that FUS undergoes nonhistone lactylation through the lactylation modification “Writer” EP300. Immunofluorescence images revealed the colocalization of FUS and EP300 in the cell nucleus (Figure 6H). Concurrently, C646 significantly reversed the changes in ferroptosis-related indicators (Figure 6I) and the expression of SLC7A11 (Figure 6J), as well as the changes in the ferroptosis-related proteins ACSL4 and GPX4 (Figure S6I). These results indicate that lactylation modified by EP300 plays a role in regulating the ferroptosis of HPASMCs caused by hypoxia.

We speculated that EP300 undergoes phase separation with ca-circSCN8A and FUS. To verify this speculation and elucidate the mechanism of nonhistone FUS lactylation modified by EP300, we first used the IUpred2 program to predict that EP300 indeed has many disordered regions required for LLPS (Figure S6J). The mCherry fusion EP300 expression vector was constructed and transfected into HPASMCs. The droplet fusion experiment (Figure 6K) and fluorescence recovery after the photobleaching experiment (Figure 6L) revealed that EP300 can undergo LLPS. Then, we used confocal microscopy to observe the formation and distribution of ca-circSCN8A, FUS, EP300, and aggregates in HPASMCs. The results revealed that three types of nuclear aggregates formed in HPASMCs induced by hypoxia, and knocking down EP300 (Figure S6K) and using the EP300 inhibitor C646 (Figure 6M) to treat hypoxic cells interrupted LLPS. Thus, the LLPS formed by ca-circSCN8A and FUS is regulated by the lactylation modification “writer” EP300.

Given that ca-circSCN8A colocalizes with FUS and EP300, we next explored whether ca-circSCN8A can induce the formation of LLPS composed of FUS. We found that in hypoxia-induced ca-circSCN8A-silenced HPASMCs, the number of LLPS aggregates was significantly reduced (Figure 6N). In addition, to determine the effect of ca-circSCN8A on the formation of FUS and EP300 droplets in vitro, we purified total RNA from HPASMCs, transcribed and cyclized ca-circSCN8A RNA in vitro, and coincubated it with FUS or EP300 without levoglucosan T500. In contrast to tRNA, total RNA can promote droplets of FUS, EP300, and a mixture of FUS and EP300. However, ca-circSCN8A induced droplets of FUS and a mixture of FUS and EP300 but failed to induce droplets of single EP300 (Figure 6O). Moreover, the longer transcript fragments of ca-circSCN8A had a stronger driving force for the droplets of FUS and EP300 (Figure 6P). These studies show that ca-circSCN8A promotes the formation of the LLPS of FUS and EP300 in a long-term and concentration-dependent manner and that EP300 is essential in this process. In summary, the above research results show that ca-circSCN8A recruits EP300 to promote nonhistone FUS lactylation-mediated LLPS under hypoxic conditions.

**Figure 6:**
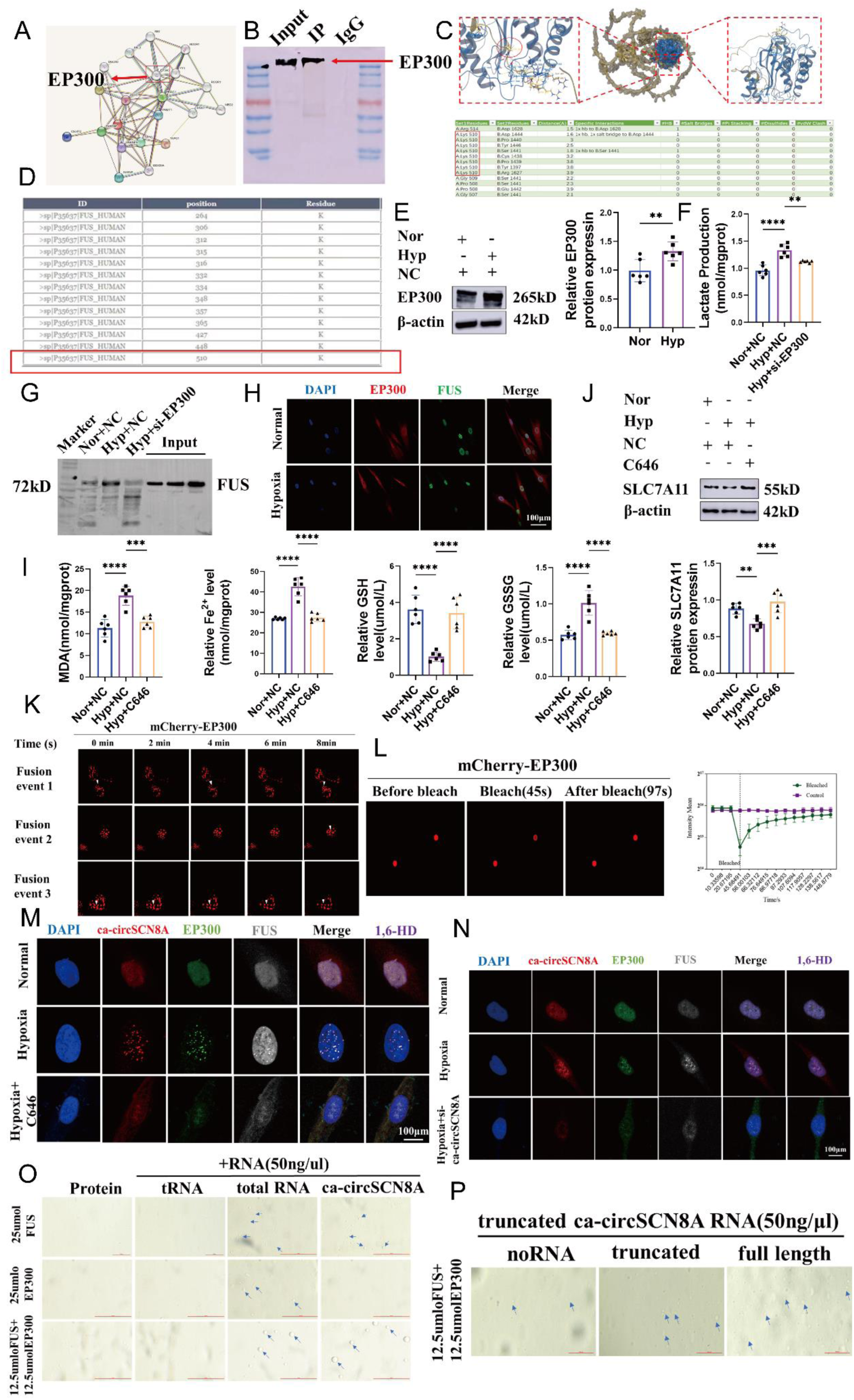
LLPS formed by ca-cicSCN8A and FUS is regulated by the lactate-modified “writer” EP300. (A) The STRING database was used to predict downstream binding proteins of FUS. (B) Co-IP experiment to verify the interaction between FUS and EP300. (C) A protein‒protein interaction simulation was used to predict the binding site between FUS and EP300. (D) Bioinformatics analysis of FUS lactate sites. (E) WB was used to measure the expression of EP300 in normoxic and hypoxic HPASMCs (n = 6). (F) A lactate assay kit was used to measure the expression levels of lactate in normal oxygen, low-oxygen, and low-oxygen si-EP00 HPASMCs (n = 6). (G) Immunoprecipitation and immunoblotting experiments were used to measure the expression levels of si-EP300 and FUS lactate under normoxia, hypoxia, and hypoxia. (H) Immunofluorescence images of the colocalization of EP300 and FUS. (I) The impact of the EP300 inhibitor C646 on the expression levels of iron death-related indicators Fe^2+^, MDA, GSH, and GSSG in HPASMCs was measured via the relevant reagent kits (n = 6). (J) WB was used to measure the effect of C646 on the expression of SLC7A11 (n=6). Droplet fusion experiments (K) and photobleaching recovery experiments (L) verified the ability of EP300 to form LLPS. (M) Observation of the formation and distribution of cicSCN8A, FUS, and EP300 LLPS nuclear condensates via confocal microscopy, as well as the effect of C646 on the formation of LLPS nuclear condensates. (N and P) Microscopic observation of the effect of ca-cicSCN8A on droplet formation in vitro. (O) Observation of the formation and distribution of ca-circSCN8A, FUS, and EP300 LLPS nuclear condensates via confocal microscopy, as well as the effect of si-ca-circSCN8A on LLPS nuclear condensates.

## DISCUSSION

Ferroptosis plays a crucial role in cardiovascular diseases.^25, 26^ However, the molecular mechanisms underlying ferroptosis in PH remain unexplored. Our study highlights the following new concepts. First, ca-circSCN8A, a novel regulatory factor, mediates HPASMC redox-dependent ferroptosis by regulating the transcription of the nonhost SLC7A11 gene. Second, ca-circSCN8A regulates SLC7A11 gene transcription by forming an R-loop structure with the nonhost gene SLC7A11 promoter. Third, the formation of an R-loop between ca-circSCN8A and the SLC7A11 promoter is driven by LLPS generated by nonhistone lactication-modified FUS in the binding of ca-circSCN8A and the SLC7A11 promoter. Our results reveal a novel mode by which ca-circSCN8A regulates gene transcription. In addition, we predicted the splicing factors of pre-mRNAs containing exons of ca-circSCN8A and found a significant association with SRSF1. We hypothesize that hypoxia may upregulate ca-circSCN8A expression through SRSF1.

Emerging evidence suggests that circRNAs can promote or inhibit ferroptosis by regulating key gene transcription and proteins after transcription.^27–30^ CircRNAs are involved in ferroptosis by regulating the metabolism of iron, reactive oxygen species (ROS), and ferroptosis-related amino acids. The relationship between circRNAs and ferroptosis therefore influences disease progression and offers novel targets for disease treatment.^31^ Through bioinformatics analysis of clinical data from PH patients, we identified a novel gene, ca-circSCN8A. Furthermore, our results revealed that ca-circSCN8A is upregulated both in vivo and in vitro. Moreover, ca-circSCN8A mediates ferroptosis in hypoxic HPASMCs by modulating the transcription of the ferroptosis-related cystine/glutamate antiporter SLC7A11, not ACSL4 or GPX4. Our findings are the first to indicate that ca-circSCN8A plays an important role in the ferroptosis of hypoxia-induced HPASMCs via the regulation of nonhost gene expression and provide a novel potential target for the treatment of PH.

During gene transcription, newly synthesized RNA forms R-loops, which play a significant role in gene transcription and DNA damage repair.^32^ The abnormal accumulation of these genes can lead to genomic instability. circRNAs have been reported to form a covalent loop^33,34^ and can regulate the transcription of their host genes by forming an R-loop.^35^ Surprisingly, our bioinformatics analysis revealed sequence complementarity between ca-circSCN8A and the SLC7A11 gene promoter. The ChIRP-qPCR results confirmed that ca-circSCN8A forms an R-loop with the SLC7A11 promoter at site 1, thereby regulating its expression and mediating ferroptosis in hypoxia-treated HPASMCs. To our knowledge, this is the first experimental confirmation that circRNAs form R-loops with nonhost genes. Our findings indicate new avenues for understanding the complex interactions and regulatory mechanisms involved in cellular biology.

To explore whether ca-circSCN8A forms an R-loop by binding to specific proteins, we used bioinformatics analysis to predict the downstream proteins of ca-circSCN8A and selected FUS because it demonstrated the highest binding affinity. Our RNA pull-down and RIP experiments further confirmed that ca-circSCN8A and FUS can bind to HPASMCs. Interestingly, our results revealed that ca-circSCN8A does not regulate the expression level of FUS, suggesting that ca-circSCN8A induces R-loop formation and is indirectly regulated by FUS. Moreover, our analysis revealed that ca-circSCN8A is significantly enriched in the nucleolar speck pathway. A previous study showed that FUS can undergo phase separation,^36, 37^ and FUS contains IDRs that recognize R-loop structures.^38^ Therefore, we speculate that ca-circSCN8A may undergo phase separation with FUS and further form an R-loop with the SLC7A11 promoter. We used confocal microscopy to observe that in the nuclei of hypoxic HPASMCs, Cy3-labeled ca-circSCN8A and FITC-labeled FUS can form aggregates. Moreover, droplet fusion and photobleaching recovery experiments confirmed that the aggregate was produced by LLPS. Notably, an inhibitor of LLPS can significantly reduce the R-loop. The above studies focused mainly on the nucleus, and the molecular mechanism of ca-circSCN8A in the cytoplasm of hypoxic HPASMCs remains unclear and will be the focus of our future investigations.

Studies have shown that the IDR of proteins regulates the formation of LLPS and the assembly of stress granules (SGs) through acetylation/deacetylation.^39^ In the present study, we used STRING to predict the downstream binding proteins of FUS and similarly selected EP300, which is an enzyme that acetylates or lactylates histones. We also found that the two proteins can bind LYS 510. Interestingly, we predicted that LYS510 is the lactylation site of FUS. Many types of posttranslational modifications can regulate LLPS,^40^ and circRNA can indirectly modulate lactylation. For example, circXRN2 can inhibit tumor growth by suppressing histone lactylation through the activation of the Hippo pathway.^41^ Therefore, we speculate that FUS forms a complex with EP300, causing the lactylation of FUS, and further binds with ca-circSCN8A to form LLPS. ca-circSCN8A is potentially capable of recruiting EP300, thereby forming a complex with FUS. This interaction could lead to the lactylation of FUS, which further results in the formation of LLPS. In hypoxia-treated HPASMCs, EP300 expression was upregulated. When we knocked down the expression of EP300, the lactylation of FUS significantly decreased, and the formation of LLPS was also significantly reduced. To our knowledge, this is the first study in which nonhistone lactylation has been demonstrated to regulate the formation of LLPS. While the result is promising, this study is limited by the inability to observe LLPS in vivo, owing to current experimental technique constraints.

In conclusion, our findings suggest that hypoxia-induced ca-circSCN8A and lactylation-mediated LLPS play crucial roles in regulating ferroptosis in PH. Hypoxia recruits lactate-modified FUS to undergo LLPS through ca-circSCN8A in the cell nucleus. The disordered region of FUS can bind to R-loops at the SLC7A11 gene promoter, causing ca-circSCN8A to accumulate in the R-loop region of the SLC7A11 gene promoter. This promotes the formation and accumulation of R-loop structures, downregulates the expression of SLC7A11, leads to an imbalance in the redox state of PASMCs, and mediates ferroptosis (Figure 7).

**Figure 7:**
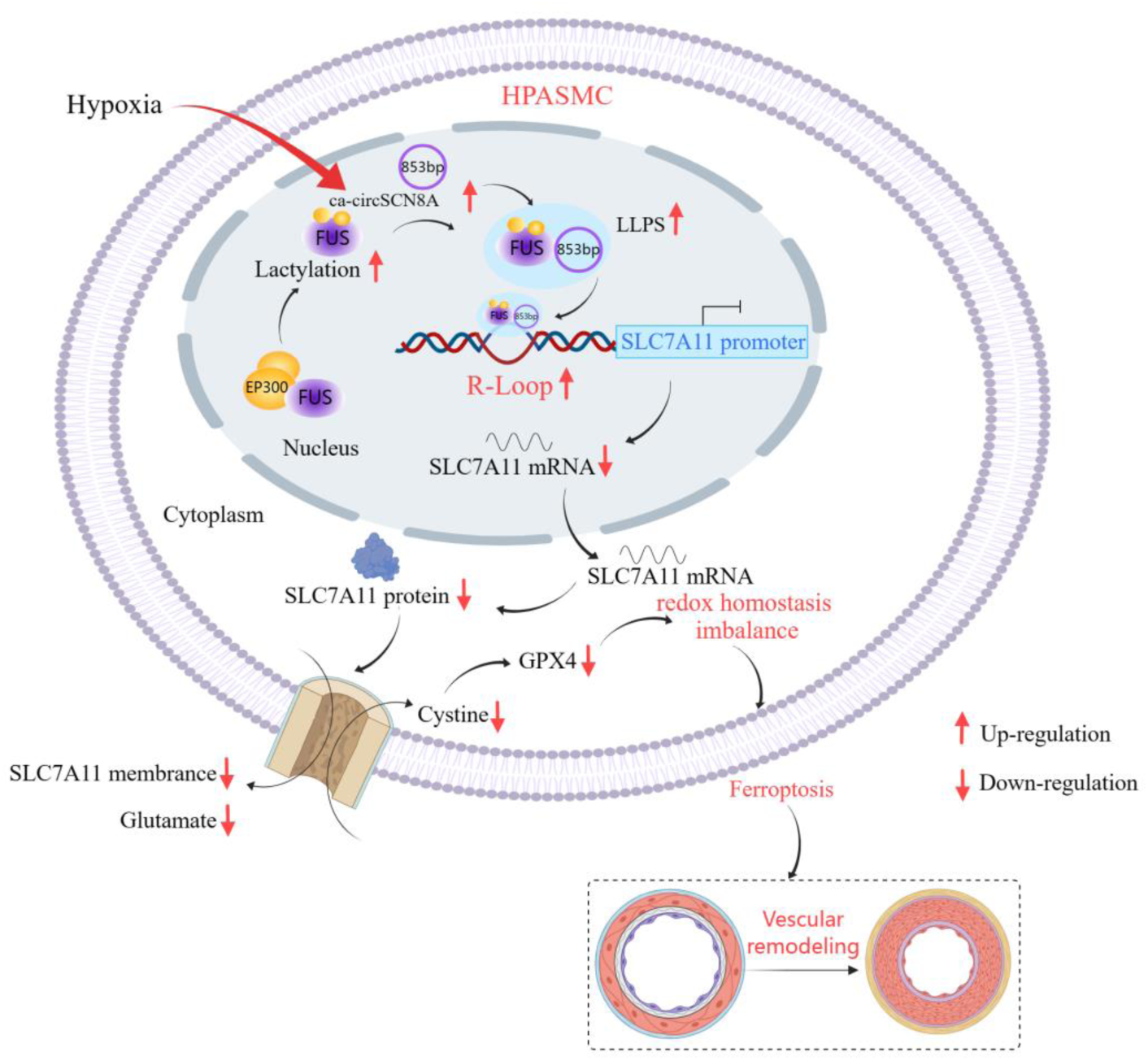
Schematic of the conclusion. Under hypoxic conditions, lactate-modified FUS is recruited via ca-cicSCN8A within the nucleus, thereby inducing LLPS. The disordered region of FUS has the ability to bind to the R-loop region of the SLC7A11 gene promoter. This binding process enriches ca-cicSCN8A in the R-loop region of the SLC7A11 gene promoter, which in turn promotes the formation and accumulation of the R-loop structure. Consequently, the expression of SLC7A11 is downregulated, leading to disruption of the redox homeostasis of PASMCs and mediating ferroptosis.

## Nonstandard Abbreviations and Acronyms

PH: pulmonary hypertension
PVR: pulmonary vascular remodeling
HPASMC: human pulmonary arterial smooth muscle cell
ca-circRNA: chromatin-associated circRNA
FRAP: fluorescence recovery after photobleaching
AAV9: adeno-associated virus serotype 9
ASO: antisense oligonucleotide
SuHX: sugen5416
SLC7A11: solute carrier family 7 member 11
GPX4: glutathione peroxidase 4
ACSL4: acyl-CoA synthetase long-chain family member 4
GSSG: glutathione
GSH: reduced glutathione
MDA: malondialdehyde
LLPS: liquid-liquid phase separation

## Acknowledgments

We thank LetPub (www.letpub.com.cn) for its linguistic assistance during the preparation of this manuscript.

## DATA AVAILABILITY

The data underlying this article will be shared on reasonable request to the corresponding author.

## Supplementary Data statement

Supplementary Data are available at NAR online.

## AUTHOR CONTRIBUTIONS

Mengnan Li: Conceptualization, Formal analysis, Methodology, Validation, Writing—original draft. Daling Zhu: Conceptualization, Formal analysis, Visualization, Writing—review & editing. Yingying Hao, Xinyue Song, Chi Zhang, Jiaqi Zhang and Huiyu Liu: Formal analysis, Methodology. Hanliang Sun: Methodology, Xiaodong Zheng, Cui Ma, Hang Yu, Lixin Zhang and Xijuan Zhao: Formal analysis.

